# Left-handedness and its genetic influences are associated with structural asymmetries mapped across the cerebral cortex in 31,864 individuals

**DOI:** 10.1101/2021.07.16.452594

**Authors:** Zhiqiang Sha, Antonietta Pepe, Dick Schijven, Amaia Carrion Castillo, James M. Roe, René Westerhausen, Marc Joliot, Simon E. Fisher, Fabrice Crivello, Clyde Francks

## Abstract

Roughly 10% of the human population is left-handed, and this rate is increased in some brain-related disorders. The neuroanatomical correlates of hand preference have remained equivocal. We re-sampled structural brain image data from 28,802 right-handers and 3,062 left-handers (UK Biobank population dataset) to a symmetrical surface template, and mapped asymmetries for each of 8,681 vertices across the cerebral cortex in each individual. Left-handers and right-handers showed average differences of surface area asymmetry within fusiform, anterior insular, anterior-middle-cingulate and precentral cortex. Meta-analyzed functional imaging data implicated these regions in executive functions and language. Polygenic disposition to left-handedness was associated with two of these regional asymmetries, and 18 loci previously linked with left-handedness by genome-wide screening showed associations with one or more of these asymmetries. Implicated genes included six encoding microtubule-related proteins: TUBB, TUBA1B, TUBB3, TUBB4A, MAP2 and NME7 – the latter is mutated in left-right reversal of the visceral organs. There were also two cortical regions where average thickness asymmetry was altered in left-handedness: on the postcentral gyrus and inferior occipital cortex, functionally annotated with hand sensorimotor and visual roles. These cortical thickness asymmetries were not heritable. Heritable surface area asymmetries of language-related regions may link the etiologies of hand preference and language, whereas non-heritable asymmetries of sensorimotor cortex may manifest as consequences of hand preference.

## Introduction

Roughly 90% of the human population is right-handed and 10% left-handed, and this strong bias is broadly consistent across cultures, ethnicities, and history (1–5). Hand-motor control is performed primarily by the contralateral brain hemisphere, such that right handedness reflects left-hemisphere specialization for hand articulation in most people (6). Behavioural precursors of hand preference are established already in the developing human fetus (7, 8), probably as a consequence of a genetically regulated program of asymmetrical brain development (9–12). Left-handedness occurs at increased frequencies in neurodevelopmental and psychiatric conditions including intellectual disability (13), autism (14), and schizophrenia (15), which indicates that altered asymmetry may sometimes be a marker of disrupted neurodevelopment – however, it is important to stress that the large majority of left-handed people experience no adverse functional effects whatsoever.

Despite decades of research, the neuroanatomical correlates of hand preference have remained uncertain. Previous magnetic resonance imaging (MRI) studies have indicated slightly altered lateralization of the morphology of sensorimotor cortex around the central sulcus as well as temporal auditory cortex in left-handed people, if they found any effects at all (16–22). Most of these studies focused on hypothesis-driven regions of interest rather than mapping across the cortex, and they used different methods of analysis and measurement. Results were conflicting, not replicated or negative, possibly due to small sample sizes; the largest number of left-handers in any of these studies was 198 (16). Cortex-wide screening using atlas-defined cortical regions has also not shown robust associations with hand preference (23, 24), even in as many as 608 left-handers vs. 7,243 right-handers. One possibility is that the neuroanatomical correlates of handedness may be too focal to be captured by common atlas-defined parcellations. To this end, vertex-wise mapping in a large population-sample may provide new insights into how hand preference relates to cerebral cortical structural asymmetry.

The most statistically robust association of handedness with brain structure reported to date reflects altered average whole-hemispheric skew (or ‘torque’) in the horizontal and vertical planes, found in 35,338 right- and 3,712 left-handed adults from the UK Biobank (25). However, an association of whole-hemispheric torque with handedness does not identify specific brain regional asymmetries linked to this behavioural trait. Associations of left-handedness have also been reported with increased functional connectivity between left and right language networks in roughly 9000 UK Biobank individuals (26), in analysis of resting-state functional MRI data. The asymmetry of intra-hemispheric functional connectivity during the resting state has also been associated with hand preference (27). The left hemisphere is dominant for language in more than 95% of the right-handed population, but in only around 70% of the left-handed population, which suggests possible developmental and evolutionary relationships between these two functional asymmetries (28–33).

In the present study, we mapped cerebral cortical structural asymmetry with respect to hand preference, using a large sample and atlas-free approach: 28,802 adult right-handers and 3,062 left-handers from the UK Biobank, measured for asymmetries of cortical surface area and thickness at each of 163,842 vertices in each hemisphere, before down-sampling the asymmetry maps to 8,681 vertices to test associations with hand preference. Vertex-wise correspondence between the left and right hemispheres was achieved through re-sampling each individual’s cortical surface model to a symmetrical template created by inter-hemispheric co-registration (34, 35). This surface-based registration approach aligns cortical folding patterns across individuals and both hemispheres.

Using these data, we first aimed to identify specific clusters of vertices where cortical surface area or thickness asymmetries differed significantly at the group level between right and left-handers, for a uniquely well-powered and precision mapping analysis of the cortical correlates of hand preference. We then used meta-analyzed functional MRI (fMRI) data to annotate the cognitive and behavioural functions of the implicated cortical regions, based on independent studies.

Handedness is a weakly but significantly heritable trait, with estimates of the heritability due to common genetic polymorphisms ranging from 1.2% to 5.9% across different large-scale studies and cohorts (36, 37). Recent large-scale genome-wide association studies have made progress on the identification of genetic influences on handedness (26, 36, 37), and separately also on brain structural asymmetry (11). However, no study has previously investigated shared genetic influences on handedness and its specific cortical structural correlates. Using genome-wide genotype data for single nucleotide polymorphisms (SNPs) in the UK Biobank individuals, we tested the heritabilities of the cortical asymmetry measures at each regional cluster associated with hand preference, and also the genetic correlations between these regional cortical asymmetries. We then tested whether polygenic disposition to left-handedness was associated with the specific cortical regional asymmetries that are associated with this behavioral trait, including the use of mediation analyses. In addition, we tested 39 individual SNPs in relation to the asymmetries of these cortical regions – these SNPs were previously implicated in left-handedness at a significant level after adjustment for multiple testing over the entire genome, in a study of 1,766,671 individuals. These analyses would reveal numerous insights into the biology of gene-brain-behavioral links involving human hand preference.

## Results

### High-resolution mapping of hand preference in relation to cerebral cortical asymmetry in 31,864 individuals

For each of 28,802 right-handed and 3,062 left-handed adult individuals from the UK Biobank, we generated a cortical surface area asymmetry map, and a cortical thickness asymmetry map, where asymmetries were ultimately measured for each of 8,681 vertices across the cortical surface using an asymmetry index: AI=(Left-Right)/((Left-Right)/2) (Fig. 1; see Materials and Methods).

**Figure 1.**
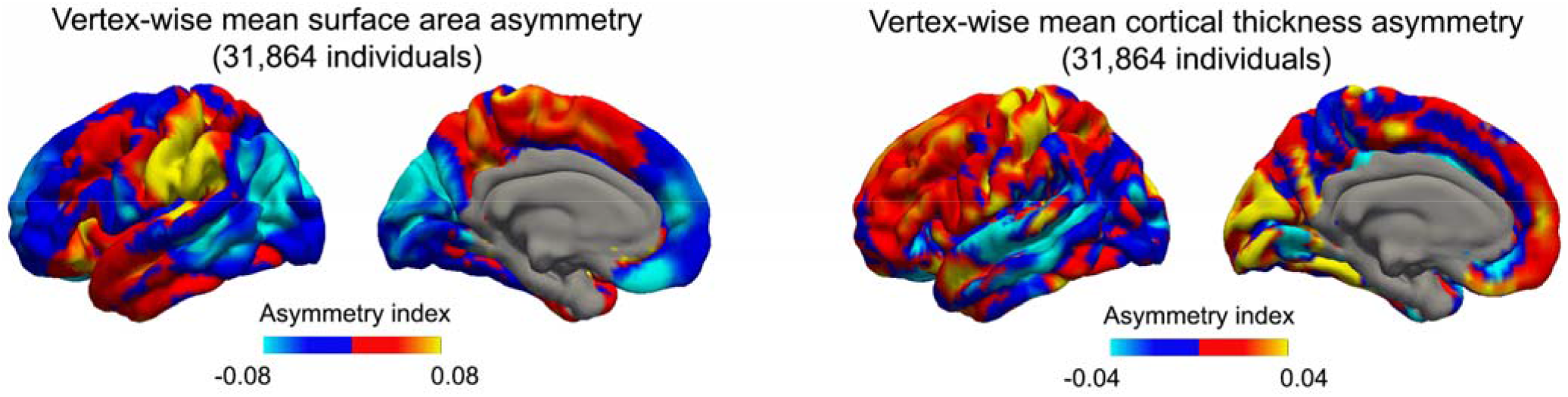
Maps of average cortical asymmetries across 31,864 adult individuals (28,802 right-handers and 3,062 left-handers). Red-orange-yellow colour indicates vertex-wise leftward (Left>Right) average asymmetry, whereas blue indicates vertex-wise rightward (Left<Right) average asymmetry. Vertex-wise correspondence between hemispheres and individuals was achieved through re-sampling each individual’s cortical model to a symmetrical template based on inter-hemispheric co-registration.

For cortical surface area asymmetries, we found eight significant clusters (cluster-level corrected p<0.05) where left-handers differed on average from right-handers: three on the anterior insula (most significant peak t-value=4.98, p-value=6.55×10^−7^), one on the precentral gyrus (peak t=4.35, p=1.34×10^−5^), three in anterior-middle-cingulate cortex (most significant cluster peak t=4.43, p=9.32×10^−6^), and one on the fusiform cortex (peak t=4.74, p=2.13×10^−6^; Fig. 2, Supplementary Figures 1-2, Supplementary Table 1). The most anterior of the insula clusters also extended onto the neighboring pars triangularis (Fig. 2).

**Figure 2.**
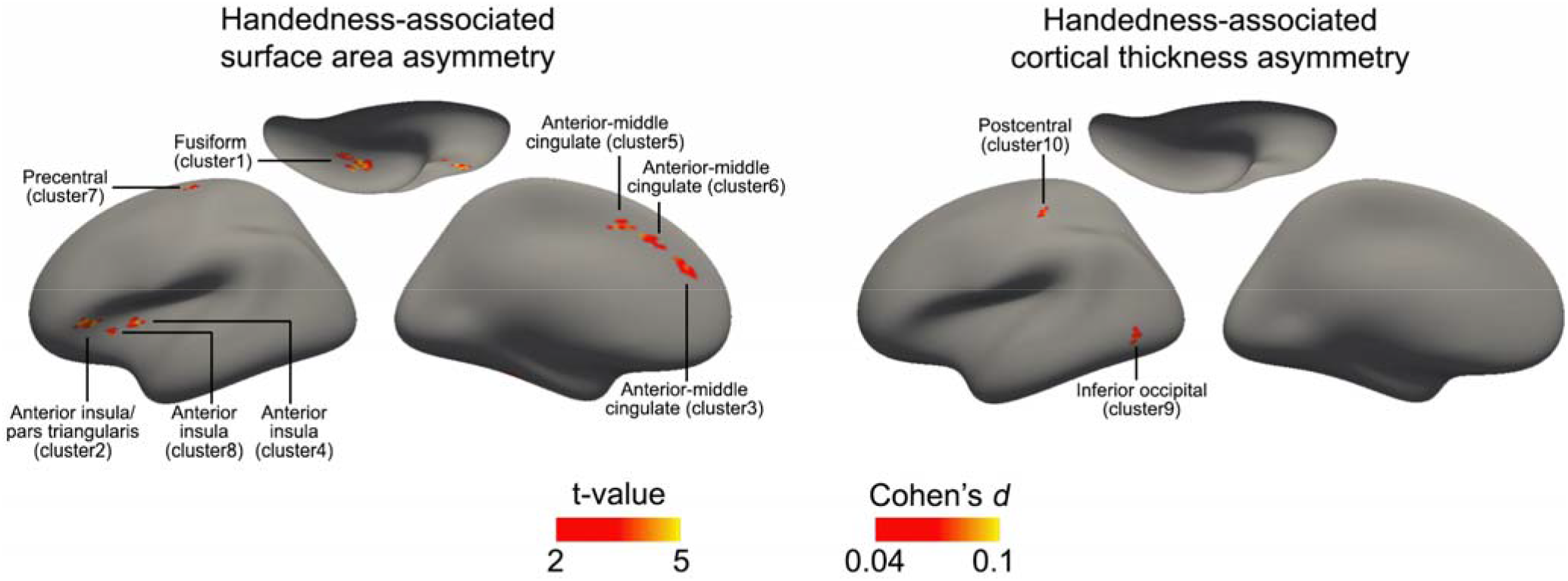
Cerebral cortical asymmetries associated with human hand preference. Significant clusters of association between hand preference and cerebral cortical structural asymmetries, for surface area (left) and cortical thickness (right). All clusters showed lower asymmetry indexes in left-handers compared with right-handers (i.e. less leftward or more rightward average asymmetries in left-handers). There were ten clusters in total that survived multiple testing correction (vertex-wise p<0.001; cluster-wise p<0.05). For all maps, the colour scale indicates both the t values and their corresponding Cohen’s *d* values.

For cortical thickness asymmetries, there were two clusters where left-handers differed on average from right-handers (cluster-level corrected p<0.05): one on the postcentral gyrus (peak t=4.05, p=4.12×10^−5^) and one on the inferior occipital gyrus (peak t=4.09, p=4.39×10^−5^; Fig. 2 and Supplementary Table 1).

Strikingly, for all surface area and thickness asymmetries within these significant clusters, left-handers showed lower average asymmetry indexes than right-handers: in other words, if a cluster had an average rightward asymmetry in right-handers, it had a stronger average rightward asymmetry in left-handers, and if a cluster had an average leftward asymmetry in right-handers, it had a weaker average leftward asymmetry in left-handers (Supplementary Figure 1, Supplementary Table 1). These data indicate that left-handedness, at the group level, is associated with a relative shift of neural resources to the dominant right hemisphere for hand motor control, within all handedness-associated cortical clusters. The unilateral hemispheric effects corresponding to the clusters with altered asymmetry in left-handers are in Supplementary Table 2.

### Functional annotation of cortical regional asymmetries linked to hand preference

We used meta-analyzed fMRI activation data from 14,371 studies, compiled within the Neurosynth database (38), to annotate our handedness-associated structural asymmetry maps with cognitive and behavioural functions (see Materials and Methods).

We found that the clusters with altered average surface area asymmetries in left-handedness are especially activated by tasks involving executive functions, language and reading, mood and pain perception (Fig. 3 and Supplementary Table 3). The language-related annotations such as ‘word, phonological, orthographic, reading, language’ were likely contributed by the anterior insula and fusiform clusters in particular (Fig. 2): clusters 2 and 8 on the anterior insula, and cluster 1 on the fusiform cortex, overlap with regions previously identified as being conjointly and left-asymmetrically activated in each of the three sentence-level language task contrasts (39). Hemispheric language dominance is known to associate with hand preference, as atypical right-hemispheric language dominance is more frequent in left-handed than right-handed people (29, 40).

**Figure 3.**
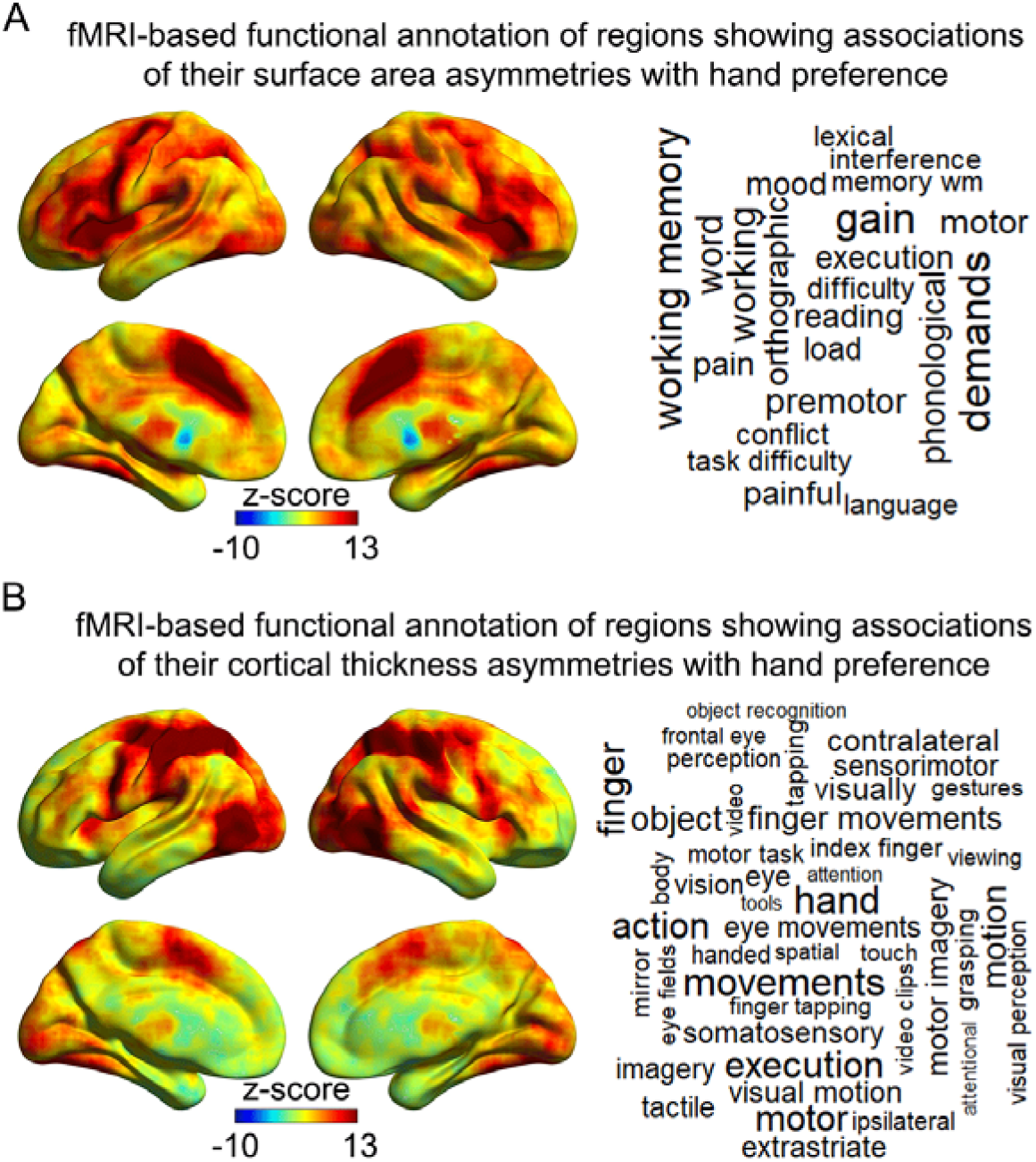
Functional annotation of regions that show altered asymmetry in left-handedness. Functional annotation based on meta-analyzed fMRI data, for clusters showing altered surface area asymmetry (A) and cortical thickness asymmetry (B) in left-handedness. Left panel: co-activation maps for the handedness-associated clusters, derived from the “decoder” function of Neurosynth (higher z-scores indicate greater co-activation). Right panel: word clouds of cognitive terms associated with the co-activation maps for the handedness-associated clusters. The font sizes of terms indicate the correlations of their meta-analytic activation maps with the co-activation maps for the handedness-associated clusters. See also Supplementary Table 3.

We found that clusters with altered cortical thickness asymmetries in left-handedness are especially activated by tasks involving hand and finger movement, as well as somatosensory and tactile tasks (Fig. 3 and Supplementary Table 3), likely contributed in particular by the postcentral gyrus cluster. The top functional terms were ‘movements, hand, execution, finger, action, motor, motion’ (Fig. 3 and Supplementary Table 3). These functional annotations based on independent fMRI data serve as a strong validation of the cortical thickness asymmetry associations with left-handedness that we observed (Fig. 3 and Supplementary Table 3). There were also numerous visual-system-related functional annotations for the regions with altered cortical thickness asymmetries in left-handedness (Supplementary Table 3), likely contributed especially by the inferior occipital cluster (Fig. 2).

The term ‘motor’ was annotated to the clusters showing altered surface area asymmetry and to those showing altered thickness asymmetry in left-handedness (Supplementary Table 3), again supporting the validity of these anatomy-handedness associations.

### Genetic disposition to left-handedness associates with specific cortical asymmetries that are altered in left-handedness

We made use of UK Biobank genome-wide genotype data to calculate the SNP-based heritabilities (41) of the cortical asymmetry measures for each separate cluster that was associated with hand preference, (Supplementary Table 4; Materials and Methods). These heritability values indicate the extent to which common genetic variation across the genome influences inter-individual variation in each of the left-handedness-associated cortical regional asymmetries. The surface area asymmetries of the fusiform cluster, the three anterior insular clusters, and one anterior-middle-cingulate cluster were weakly but significantly heritable after false discovery rate (FDR) correction at 0.05, with pointwise heritability estimates ranging from 3.1% to 6.3%, (Supplementary Table 4). The surface area asymmetry of the precentral gyrus cluster was not significantly heritable (pointwise heritability 0.0, p=0.5). Likewise, the cortical thickness asymmetries associated with left-handedness were not significantly heritable, with pointwise estimates no greater than 1.4% (lowest p=0.10).

Genetic correlation analysis indicated that the surface area asymmetries of the three anterior insula clusters were influenced by largely the same genetic variants across the genome, with significant pairwise genetic correlations from 0.85 to 1.0, whereas other pairwise genetic correlations between heritable regional asymmetries were weaker (see phenotypic and genetic correlations in Supplementary Tables 5 and 6).

From an independent set of 272,673 right-handed and 33,704 left-handed individuals in the UK Biobank, who were not overlapping and unrelated to those with brain image data, we derived effect sizes for the association of each SNP across the genome with hand preference (see Materials and Methods). We then applied these effect size estimates, in mass combination, to the 28,802 right-handed and 3,062 left-handed individuals having genetic and brain imaging data for the present study, in order to quantify their polygenic dispositions to left-handedness, using the PRS-CS toolbox (42). Polygenic disposition to left-handedness was significantly associated with hand preference in this sample as expected (logistic regression beta=0.12, p=6.63×10^−5^), as well as cerebral cortical surface area asymmetry within the fusiform cluster (r=−0.02, p=0.003), and anterior insula clusters 2 (r=−0.02, p=6.59×10^−4^) and 4 (r=−0.02, p=9.01×10^−5^) after FDR correction at p<0.05 (Fig. 4 and Supplementary Table 7). Specifically, higher polygenic disposition to left-handedness was associated with increased average rightward asymmetry in the fusiform cluster, and decreased average leftward asymmetry in the anterior insula clusters (Fig. 4), consistent with the associations of these regional surface area asymmetries with the left-handedness trait itself. The polygenic disposition to left-handedness was not associated with any other regional cortical asymmetries linked to left-handedness, including the precentral surface area asymmetry cluster, and the postcentral thickness asymmetry cluster (Fig. 4 and Supplementary Table 7).

**Figure 4.**
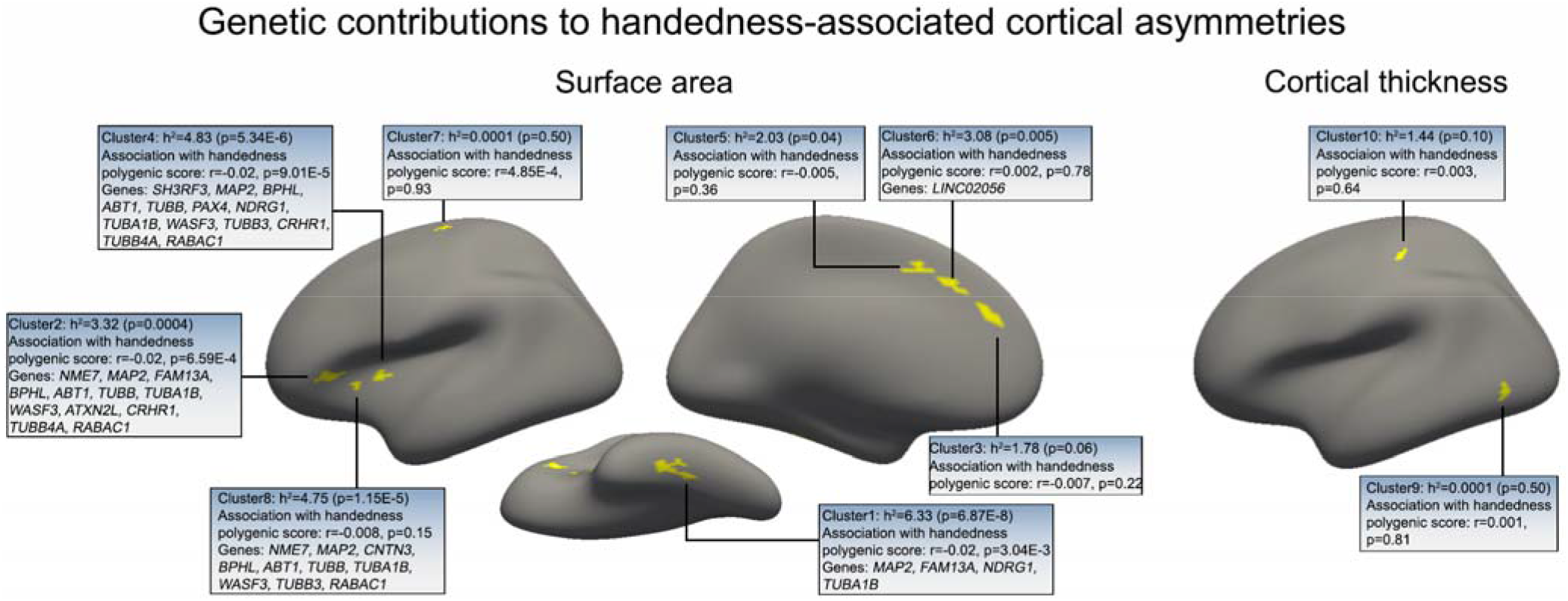
Genetic influences on hand preference impact cortical asymmetries linked to hand preference. Five clusters of handedness-associated cortical surface area asymmetry showed significant heritabilities after FDR correction. i.e., three clusters in the anterior insular cortex, one in the fusiform cortex and one in the anterior-middle-cingulate cortex. Other handedness-associated asymmetry clusters were not significantly heritable, including the pre- and postcentral clusters. The associations of cortical asymmetries with polygenic disposition to left-handedness are indicated, as well as individual handedness-associated genes implicated in their variabilities.

Separately for the fusiform cluster and each anterior insula cluster that showed association with polygenic disposition to left-handedness, we applied mediation analyses according to two possible causal models: polygenic disposition>asymmetry>handedness, and polygenic disposition>handedness>asymmetry (see Materials and Methods). All of the models showed evidence of both direct and mediated effects (Supplementary Tables 8 and 9).

### Individual genomic loci associated with hand preference and its cerebral cortical correlates

A previous genome-wide association study, based on 1,766,671 individuals, found 41 individual genomic loci that were significantly associated with left-handedness after multiple testing correction over the whole genome (37). In the 28,802 right-handed and 3,062 left-handed individuals with both genetic and brain imaging data in the present study, we found that 18 of these genomic loci were associated with at least one of the specific cortical regional surface area asymmetries that are linked to left-handedness, after FDR correction at 0.05 (Supplementary Table 10). In particular, 17 out of these 18 SNPs were associated with surface area asymmetry of either the fusiform cluster or at least one of the anterior insula clusters (Supplementary Table 10).

The protein-coding genes located nearest to these SNPs in the genome include four that encode tubulin components of microtubules, TUBB, TUBA1B, TUBB3 and TUBB4A, as well as microtubule-associated proteins MAP2 and NME7. These findings are in line with another recent study which implicated microtubule-related genes in brain structural asymmetry, in that case using atlas-based regional definitions across the whole brain, but without considering handedness (11). Recessive mutations in *NME7* cause *situs inversus totalis*, a rare condition in which the visceral organs (heart, liver etc.) are reversed in their orientation on the left-right axis (43). Other genes implicated by the present study to affect both handedness and its specific cerebral cortical correlates include *CNTN3*, which mediates cell surface interactions during nervous system development (44), and *ATXN2L*, implicated in macrocephaly by *de novo* mutation (45).

## Discussion

### Cerebral cortical structural asymmetry with respect to hand preference

In this study we mapped cerebral cortical correlates of hand preference in 28,802 right-handed and 3,062 left-handed adults using an atlas-free, cortex-wide approach. This was by far the largest study of this question to date, and brings clarity to a long-standing issue in human neuroscience, where previous studies had reported contradictory, equivocal and/or negative results (see Introduction). The vertex-level mapping, made possible through re-sampling to a symmetrical template constructed through inter-hemispheric registration – meant that we were able to detect and delineate regional cortical correlates of hand preference that would likely remain concealed with atlas-based parcellation. We found that, at the group level, left-handers had an average less leftward/more rightward shift of specific regional cortical surface area or thickness asymmetry than right-handers, distributed in several cortical regions – fusiform, anterior insula (partly overlapping with the pars triangularis), precentral, postcentral, anterior-middle-cingulate and inferior occipital cortex. For all of these regions, the average alterations were consistent with a relative shift of neural resources to the right hemisphere that controls the dominant hand in left-handers. Together, the implicated regions suggest a distributed neural network associated with controlling articulation of the dominant hand.

Functional annotation of the regions with altered average surface area asymmetries in left-handers, based on independent fMRI data from 14,371 meta-analyzed studies, pointed to their involvements in executive functions including working memory, as well as language and reading, mood and pain perception. Regarding language, previous studies have found an increase in atypical right-hemisphere language dominance among left-handed people (29, 40), even while most left-handers have typical left-hemisphere language dominance (see Introduction). We found an association of hand preference with surface area asymmetry within the anterior insular cortex/pars triangularis, corresponding to core regions of the language network that show left-lateralized activation during sentence-level language tasks (39). The anterior insula has been suggested to be especially important for the establishment of functional language lateralization in development, through integrating across social, emotional and attentional systems during early language acquisition (46). There were no significant associations of hand preference with asymmetries of superior temporal or middle temporal cortex, which are also core language network regions (39). Thus hand preference apparently relates to cortical structural asymmetry of frontal language regions in particular, rather than other language network regions that are closer to primary auditory cortex of the temporal lobe.

fMRI-based annotation of the cortical regions with altered average thickness asymmetries in left-handers confirmed their involvements in hand sensorimotor functions, with top annotations of ‘movements, hand, execution, finger’ and so on. This functional annotation based on independent data provides a strong confirmation of the validity of these group-level cortical asymmetry alterations in left-handedness. As noted in the introduction, some previous studies targeted the central sulcus as a region of interest and reported altered cortical anatomy there in left-handers, but based on small samples and variable methods and measures, with conflicting results, such that findings have remained uncertain. Here we have definitively implicated a specific region of the postcentral gyrus through cortex-wide, atlas-free mapping in a large sample, and shown that cortical thickness asymmetry rather than surface area asymmetry is the feature most correlated with hand preference in this region. In contrast, we found that a specific region of precentral ‘premotor’ cortex showed altered average surface area asymmetry in left-handers.

The regions that showed altered average cortical thickness asymmetry in left-handers also received numerous vision-related annotations based on fMRI data, such as ‘motor imagery, visual motion, eye movements, visual perception, eye fields’ and so on. These annotations were likely driven by the inferior occipital cluster. Left-handers have been reported to show more variable hemispheric involvement than right-handers in visual half-field tasks, which can index lateralized contributions to visual attentional tasks (47). The need for hand-eye coordination during complex manual tasks may be a specific driver of this relationship. Interestingly, the inferior occipital cluster also lies within a language network region that shows left-lateralized activation during sentence-level tasks presented in different modalities (39).

### Shared genetic influences on hand preference and its specific cerebral cortical correlates

We found that polygenic disposition to left-handedness was associated with specific handedness-linked asymmetries of surface area in the fusiform and anterior insular cortex. The directions of effect were as expected – higher polygenic disposition to left-handedness associated with reduced average leftward or increased average rightward surface area asymmetry in these cortical regions, the same as for left-handedness itself. Notably, the regional cortical thickness asymmetries associated with hand preference, i.e., within the postcentral gyrus and inferior occipital cortex, were not significantly heritable and showed no associations with polygenic disposition to left-handedness. Neither did the surface area asymmetry of the precentral gyrus cluster. Thus some specific handedness-related cortical asymmetries share genetic influences with hand preference, while others apparently do not, at least to extents that could be detected in the current sample size.

In the adult cross-sectional data of the UK Biobank it is not possible to unambiguously establish cause-effect relations. Mediation models were compatible with fusiform and anterior insula surface area asymmetries partly mediating the effect of polygenic disposition on hand preference, but also with hand preference partly mediating the effect of polygenic disposition on these brain asymmetries. Both models are plausible, as brain anatomy may in principle be both causal to behavioural outcomes and shaped plastically by repeated behaviours, or indeed merely correlated due to shared underlying factors (48). However, the cortical asymmetries that share genetic influences with hand preference seem more likely to be involved in the etiology of hand preference, especially as some of the individual genes involved may point to potential mechanisms of establishing the brain’s left-right axis in early development (see below). Furthermore, structural asymmetry of the anterior insula is already detectable from middle to late gestational age (49).

In contrast, non-heritable correlates of hand preference in the adult cortex may be more likely to reflect plastic adaptation in left-handedness, after the preference is already established developmentally. Left-handedness has shown only limited associations with the environmental and early life factors studied so far (50, 51), but is also only weakly to moderately heritable (36, 37, 52). This suggests that left-handedness arises largely from random (non-heritable) early developmental variation (53), occurring around a genetically-regulated program that is biased towards a right-handed outcome. In other words, the genetically regulated program of asymmetrical brain development appears to give rise to right-handedness with roughly 90% probability during development, and left-handedness with roughly 10% probability, while both inter-individual genetic and environmental variation seem to have generally small impacts on these probabilities. In this way, left-handedness can have non-heritable cortical correlates without necessarily invoking environmental causes.

This study pinpointed, for the first time, individual genomic loci which associate with both hand preference and some of its specific cortical structural correlates. The involvement of microtubule-related genes in human handedness and brain asymmetry suggests a possible mechanism whereby left-right asymmetry of the brain arises in the embryo from cellular chirality, i.e. an asymmetrical twisting of cellular shape, specified by the cytoskeleton (11). Such mechanisms have been described with respect to other organs of other invertebrate and vertebrate species, and can be organ-intrinsic, i.e. arise independently of other organs or systems (54–60). This seems especially pertinent to the brain, as left-hemisphere dominances for hand preference and language do not usually reverse in people with *situs inversus totalis* i.e. reversal of the visceral organs on the left-right axis, when caused by genetic mutations which affect primary ciliary components (61–64). Primary cilia play an important role in the establishment of visceral asymmetry, where the molecular chirality of the axoneme (microtubule-based central strand of the cilium) results in unidirectional ciliary motion within the embryonic node (65, 66).

In this context, it is especially notable that our data implicated *NME7* in affecting both hand preference and handedness-associated cortical surface area asymmetries. Recessive mutations in this gene cause *situs inversus totalis*, but without primary ciliary dyskinesia, i.e., ciliary function remains largely or wholly unaffected even while the visceral organs become reversed (43). The functions of this gene are not well understood, but it is known to associate with γ-tubulin and the ciliary axoneme (43) – thus it has potential functions that involve the cilium but also the microtubule cytoskeleton more generally. *NME7* provides a possible genetic link between brain and body asymmetry, via as-yet-unidentified mechanisms which do not necessarily involve ciliary dysfunction and the nodal pathway of visceral laterality formation (67). All of the genes implicated here to affect both hand preference and its cerebral cortical correlates should be studied with gene-functional approaches, such as in cellular models and knockout mice, to understand their roles in affecting brain functional and structural laterality.

### Limitations

Vertex-wise mapping of asymmetry assumes that high-resolution correspondence can be achieved between contralateral vertices in the two hemispheres. For the most part the method appears to have worked well, insofar as it distinguished some regions of association with hand preference that make sense on *a priori* grounds, such as the sensorimotor cortex of pre- and post-central sulcus, and language-related regions. However, we found it necessary to exclude a region surrounding the boundary with subcortical structures at the center of the brain (affecting entorhinal, parahippocampal and cingulate regions; Supplementary Figure 3), where extreme vertex-wise asymmetry values were present in most individuals (see Materials and Methods). This may indicate an artifact that involves extreme warping for this region, to fit the symmetrical template.

The limitation of the UK Biobank cross-sectional data for making causal inferences from associations was already mentioned above. In addition, the UK Biobank is a volunteer cohort of middle-aged to older adults, and therefore not fully representative of the population, with generally superior health and socioeconomic circumstances for this age range, and greater female participation (68). Higher risk for psychiatric disorders has been suggested to reduce participation in cohort studies (69), and we have previously found evidence for this in the UK Biobank (70).

In this study we linked genetic, brain structural and behavioural data, as well as making use of independent, meta-analyzed fMRI data to understand the functions of the implicated sets of cortical regions. Future studies of hand preference in large datasets may seek to integrate more aspects of brain structure and function from the same individuals into the analyses, including potentially resting-state fMRI and diffusion tensor imaging data on white matter structural connectivity.

The assessment of hand preference was by a simple questionnaire, where individuals could choose between four options: right-handed, left-handed, use both hands equally, or prefer not to answer. We found the ambidextrous category to be relatively unreliable over repeat visits (Materials and Methods), which led us to use a binary left/right variable for the purposes of this study. Simple assessments such as this have been shown to capture the inherent dichotomy in hand preference that is also revealed by more quantitative, multi-item questionnaires (71). However, brain-behavior association results may have been different if using performance-based hand skill measures, or semi-quantitative multi-item handedness ratings.

This study examined average group differences in cerebral cortical asymmetry between left- and right-handers. Future studies may consider the possibility that left-handedness associates with increased variability in terms of brain structure and function.

### Summary

This study used a symmetrical surface template to achieve atlas-free mapping of cerebral cortical asymmetry with respect to hand preference in an unprecedented sample size. Detailed and statistically reliable maps of the cortical correlates of human hand preference were produced, resolving a long-standing issue in human neuroscience. The implicated regions are especially involved in executive, language, motor and visual functions. Across all of the regions, the data uniformly indicated a less leftward/more rightward shift of neural resources in left-handers compared to right-handers, consistent with right-hemispheric control of the left hand. Hand preference was genetically associated with the asymmetries of frontal and fusiform language network regions, supporting developmental and evolutionary links between hand preference and language. Various other regional asymmetries associated with hand preference were not heritable, and may reflect plastic changes in support of hand preference, including those of primary sensorimotor cortex. Specific genes found to affect both hand preference and its cerebral cortical correlates include microtubule-related genes, one of which – *NME7* – provides a potential mechanistic link between brain and visceral asymmetries which may not involve impaired ciliary function. This study provides an example of how linking across human genetics, brain structure and behavior in large datasets can provide tangible new knowledge, as well as mechanistic and developmental hypotheses for future research on typical and atypical human brain function.

## Materials and Methods

### Participants

This study was conducted under UK Biobank application 16066, with Clyde Francks as principal investigator. The UK Biobank is a general adult population cohort (72, 73). The UK Biobank received ethical approval from the National Research Ethics Service Committee North West-Haydock (reference 11/NW/0382), and all of their procedures were performed in accordance with the World Medical Association guidelines. Informed consent was obtained for all participants.

#### Imaging-genetics dataset

We used the UK Biobank neuroimaging data (74, 75) released in February 2020, together with the genotype data from the same participants. To achieve a sufficiently high degree of homogeneity for the genetic analyses, we restricted the sample to participants with ‘white British ancestry’ defined by Bycroft et al. (‘in.white.British.ancestry.subset’) (72). We excluded subjects with a mismatch of their self-reported and genetically inferred sex, with putative sex chromosome aneuploidies, or who were outliers based on heterozygosity (principle component corrected heterozygosity >0.19) and genotype missingness (missing rate >0.05) (72). We randomly excluded one subject from each pair with a kinship coefficient >0.0442, as defined by the UK Biobank (72).

Handedness was assessed by a touchscreen question with four choices: right-handed, left-handed, use both right and left hands equally, prefer not to answer (UK Biobank field: 1707). We previously found that those indicating mixed handedness (roughly 2% of all individuals) had a high rate (41%) of changing their answers over repeat visits to the assessment centres, whereas those indicating either right-handedness or left-handedness at their first visit had consistency rates >97% on subsequent visits (36, 76). We therefore excluded mixed handers and focused only on those indicating either right– or left-handedness during their first visit.

These steps resulted in a final sample of 31,864 participants, comprising 28,802 right-handers and 3,062 left-handers. The age range of these participants was from 45 to 81 years (mean 63.75), and 15,064 were male, 16,800 female.

#### Genome-wide screening dataset to obtain SNP-wise effect sizes for association with hand preference

These UK Biobank individuals had genetic and handedness data, but not brain image data. We applied the same genetic and handedness exclusion criteria as above, with the additional requirement that none of these individuals should have a kinship coefficient >0.0442 with any of the individuals in the imaging-genetic data set. This yielded 272,673 right-handed and 33,704 left-handed participants, not overlapping and unrelated to the set with brain imaging data.

### MRI processing

Brain image data were derived from T1-weighted MRI scans (Siemens Skyra 3 Tesla MRI with 32-channel radio frequency receive head coil). We started from the Freesurfer 6.0 (77) ‘recon-all’ cortical reconstructions generated by the UK Biobank imaging team (UK Biobank data-field 20263, first imaging visit) – but did not make use of image-derived-phenotypes released by that team. Rather, the cortical thickness and surface area maps of both hemispheres of each individual were re-sampled into a standard symmetrical space created using inter-hemispheric co-registration (fsaverage_sym) (34), with 163,842 vertices per hemisphere (Supplementary Figure 4). This process required roughly 10 weeks of processing on 12 cluster server nodes running in parallel, and precisely aligned morphometry features between the two hemispheres to achieve vertex-wise correspondence. Separately for surface area and thickness maps, asymmetry for each left-right matched pair of vertices, in each participant, was calculated according to the asymmetry index AI=(left-right)/((left+right)/2). The asymmetry maps for surface area and thickness of each individual were then down-sampled to the left-hemisphere of template ‘fsaverage5’ (10,242 vertices per hemisphere; Supplementary Figure 4).

We noticed that a majority of individuals had extreme values of asymmetry in a region wrapping around the boundary with subcortical structures near the center of the brain (Supplementary Figure 3), which likely arises from a registration artifact. For this region we excluded any vertices from all individuals when they had surface area |AI|>1 in at least 1000 (~3%) individuals (Supplementary Figure 3) (i.e., excluded from subsequent analyses of both surface area and cortical thickness asymmetries). This left 8,681 vertex-wise asymmetry measures per individual, spanning the majority of the cerebral cortex (Fig. 1).

### Mapping cortical asymmetry correlates of hand preference

For each vertex, brain asymmetry differences between left- and right-handers were examined by two-sample t-tests, separately for surface area and cortical thickness, while controlling for continuous covariates: age when attended assessment center (fields 21003-2.0), quadratic age (age – mean_age)^2^, scanner position parameters (X, Y and Z: fields 25756-2.0, 25757-2.0 and 25758-2.0), T1 signal-to-noise ratio (field 25734-2.0), T1 contrast-to-noise ratio (field 25735-2.0), together with two categorical covariates: assessment center (field 54-2.0) and sex (field 31-0.0). Significant clusters were determined by the random field theory method at a cluster-corrected level of p<0.05 (clusters forming with vertex-level p<0.001) to minimize the likelihood of false positive results, using the SurfStat toolbox (https://www.math.mcgill.ca/keith/surfstat/). As a post-hoc analysis, we also examined the unilateral effects corresponding to clusters with altered asymmetry in left-handers. Specifically, for each hemisphere and individual, we calculated the mean unilateral value across all vertices within each significant cluster, and compared the differences between left and right-handers by two sample t-tests, while controlling for the same covariates mentioned above.

### Functional annotation of cortical regions showing altered asymmetry in left-handers

We made use of the Neurosynth database (38), a platform for large-scale automated synthesis of task-based fMRI data. This database defines brain-wide activation maps corresponding to specific cognitive or behavioural task terms, using meta-analyzed functional activation maps from studies referring to those terms (38). At the time of accessing the database it included 1307 maps, representing activation patterns corresponding to 1307 cognitive or behavioural terms, from 14,371 human neuroimaging studies. We created one bilateral mask in MNI152 standard space by labelling all clusters showing altered average surface area asymmetry in left-handers, and another bilateral mask in MNI standard space that comprised all clusters showing altered average cortical thickness asymmetry in left-handers. This was achieved through applying the ‘mri_surf2vol’ function in Freesurfer to the down-sampled, handedness-associated asymmetry cluster maps (defined in left hemisphere ‘fsaverage5’ surface space), then identifying the mirrored voxels (based on the midline of left and right hemisphere) corresponding to these clusters in the right of the symmetrical standard space, to define masks with bilaterally labelled voxels. These masks were then used separately as input to identify relevant cognitive and behavioral terms through the ‘decoder’ function of the Neurosynth database. This tool first creates a co-activation map for each input mask, based on all activation data in the database, and then tests each term-specific activation map for spatial correlation with the co-activation map. Cognitive terms with correlations >0.2 were visualized on a word-cloud plot, while excluding anatomical terms, non-specific terms (e.g. ‘Tasks’), and one from each pair of virtually duplicated terms (such as ‘Words’ and ‘Word’). This tool does not employ inferential statistical testing, but rather ranks cognitive and behavioural terms according to the correlations of their meta-analyzed activation maps with the co-activation map corresponding to a user-defined input mask.

### Heritabilities and genetic correlations of cortical asymmetries associated with hand preference

From the imputed SNP genotype data released by the UK Biobank (March 2018), 9,516,135 autosomal variants with minor allele frequencies >1%, INFO (imputation quality) score >0.7 and Hardy-Weinberg equilibrium p>1×10^−7^ were used to build a genetic relationship matrix using GCTA (78) (version 1.93.0beta). Specifically for analyses using GCTA, we further excluded one random participant from each pair having a kinship coefficient higher than 0.025 based on the calculated genetic relationship matrix (this analysis is particularly sensitive to higher levels of relatedness), resulting in 29,973 participants for this particular analysis (27,083 right-handed and 2,890 left-handed). Genome-based restricted maximum likelihood analyses, using GCTA (78), were used to estimate the SNP-based heritability for each cluster that had shown altered asymmetry in left-handedness (using the mean asymmetry index across vertices within each cluster), controlling for the above-mentioned covariates (i.e., age, (age-mean_age)^2^, scanner position parameters, T1 signal-to-noise ratio, T1 contrast-to-noise ratio, assessment center and sex), plus an additional binary covariate for genotyping array, and the first ten genetic principle components capturing population genetic diversity (UKB field ID: 22009-0.1~22009-0.10) (72), and applying FDR 0.05 across the ten clusters. Bivariate GREML analysis (79) was used to estimate genetic correlations between the pairs of significantly heritable cluster-wise asymmetries.

### Polygenic disposition to left-handedness

In the 272,673 right-handed and 33,704 left-handers who were not overlapping and unrelated to those with brain image data (see above), we again excluded genetic variants with minor allele frequencies <1%, INFO score ≤0.7, and Hardy-Weinberg equilibrium p-value ≤10^−7^. We then carried out an autosome-wide association scan for right-versus left-handedness, under an additive genetic model, with covariates for year of birth, sex, genotyping array, and the first ten genetic principle components capturing population genetic diversity, using BGENIE (v1.2) (72). SNP-wise effects on left-handedness disposition from this sample were then applied in mass combination to the 28,802 right-handed and 3,062 left-handers with post-QC imaging and genetic data (see above), to estimate their polygenic dispositions to left-handedness, using PRS-CS (42). This method uses a high-dimensional Bayesian regression framework to estimate posterior effect sizes of included SNPs, and has shown superior prediction statistics compared to approaches based on clumping by linkage-disequilibrium and P-value thresholding (80). In total, effect sizes of 1,103,636 SNPs spanning the autosomes were used, based also on their presence in the 1000 Genomes project European-descent dataset, from where PRS-CS obtains its linkage disequilibrium information (81). We used default parameters and the recommended global effect size shrinkage parameter phi=0.01. The handedness polygenic scores in the imaging-genetic dataset were z-standardized for subsequent analyses (Supplementary Figure 5).

We first tested whether polygenic disposition to left-handedness was associated with hand preference in the 28,802 right-handed and 3,062 left-handers, using logistic regression with hand preference as the outcome variable, and covariates of age, (age – mean_age)^2^, genotyping array, the first ten genetic principle components (see above), and sex.

We then extracted the mean asymmetry index over all vertices within each separate cluster that showed association of its asymmetry with left-handedness (eight clusters for surface area asymmetry, two for thickness asymmetry). Separately for each cluster’s mean asymmetry, we performed Pearson correlation analysis with polygenic disposition to left-handedness across individuals, while controlling for the same covariates as in the SNP-based heritability analysis (above), and applying FDR correction at 0.05 across the ten clusters tested.

For the asymmetry clusters that were significantly associated with polygenic disposition to left-handedness, we then applied the ‘mediation’ toolbox in R https://www.rdocumentation.org/packages/mediation/versions/4.5.0 using the same covariates and bootstrapping 1000 times, in order to test causal mediation models framed with respect to either polygenic disposition>asymmetry>handedness or polygenic disposition>handedness>asymmetry, separately for each cluster’s mean asymmetry.

### Imaging genetic analysis of individual loci associated with left-handedness

A previous genome-wide association scan for left-versus-right handedness, carried out in 1,766,671 individuals, reported 41 individually significant genomic loci after correction for multiple testing across the whole genome (37). We found genotype data for 38 of these SNPs in the 28,802 right-handed and 3,062 left-handers of the imaging-genetic sample in the present study, and were able to identify a proxy SNP for one more of these lead SNPs (rs11265393 as a proxy for rs66513715, with linkage disequilibrium r-squared of 1 in European-descent data according to LDproxy (82)). We tested each of these 39 SNPs for association with each of the significantly heritable, cortical cluster-wise asymmetries (averaged over all vertices separately within each cluster) that showed alterations in left-handers, using an additive model in BGENIE (v1.2) (72) with the same covariates as the heritability analysis (above). All 39 SNPs had imputation quality (INFO) scores > 0.7. We applied FDR correction at 0.05 to control multiple testing over all of these SNP-cluster combinations.

Note that the previous genome-wide association study of left-versus-right handedness based on 1,766,671 individuals (37) included the UK Biobank participants, and therefore also included the participants with brain image data, although the brain image data were not used in that study. The sample overlap meant that we could not use those genome-wide, SNP-wise effect size estimates to calculate polygenic disposition to left-handedness in the current study, as the genome-wide screening and target datasets must not overlap for polygenic scoring analysis. In addition, the SNP-wise effect sizes have not been made publicly available from the GWAS of 1,766,671 individuals (37).

## Acknowledgements

This research was funded by the Max Planck Society (Germany) and grants from the Netherlands Organization for Scientific Research (NWO) (054-15-101) and French National Research Agency (ANR, grant No. 15-HBPR-0001-03) as part of the FLAG-ERA consortium project ‘MULTI-LATERAL’, a Partner Project to the European Union’s Flagship Human Brain Project. This research was conducted using the UK Biobank resource under application no. 16066, with C.F. as the principal applicant. Our study made use of data generated by an image-processing pipeline developed and run on behalf of UK Biobank. The funders had no role in study design, data collection and analysis, the decision to publish or preparation of the manuscript. Many thanks to Nathalie Tzourio-Mazoyer and Bernard Mazoyer for their founding roles in the collaborative work that led to this study.

## Contributions

Z.S.: Conceptualization, methodology, analysis, visualization, original draft writing. D.S.: Methodology, analysis, review & editing. C.F.: Conceptualization, methodology, direction, supervision, original draft writing, review & editing. A.P., A.C.C., J.M.R., R.W., M.J., S.E.F. and F.C.: Conceptualization, review and editing the manuscript.

## Competing interests

The authors report no competing interests.

## Data Availability

The primary data used in this study are available via the UK Biobank website https://www.ukbiobank.ac.uk. Other publicly available data sources and applications are cited in the manuscript.

## Notes

### Competing Interest Statement

The authors have declared no competing interest.

